# Considerations for the design of vaccine efficacy trials during public health emergencies

**DOI:** 10.1101/261875

**Authors:** Natalie E. Dean, Pierre-Stéphane Gsell, Ron Brookmeyer, Victor De Gruttola, Christl A. Donnelly, M. Elizabeth Halloran, Momodou Jasseh, Martha Nason, Ximena Riveros, Conall Watson, Ana Maria Henao-Restrepo, Ira M. Longini

**Affiliations:** Department of Biostatistics, University of Florida, Gainesville, FL, USA.; World Health Organization, Geneva, Switzerland.; Department of Biostatistics, University of California, Los Angeles, CA, USA.; Department of Biostatistics, Harvard University, Boston, MA, USA.; MRC Centre for Outbreak Analysis and Modelling, Department of Infectious Disease Epidemiology, School of Public Health, Imperial College London, London, UK.; Vaccine and Infectious Disease Division, Fred Hutchinson Cancer Research Center, Seattle, WA, USA.; Department of Biostatistics, University of Washington, Seattle, WA, USA.; Medical Research Council, Banjul, The Gambia.; Biostatistics Research Branch, National Institute of Allergy and Infectious Diseases, Rockville, MD, USA.; Department of Infectious Disease Epidemiology, London School of Hygiene & Tropical Medicine, London, UK.

## Abstract

Public Health Emergencies (PHEs) provide a complex and challenging environment for vaccine evaluation. Under the R&D Blueprint Plan of Action, the World Health Organization (WHO) has convened a group of experts to agree on standard procedures to rapidly evaluate experimental vaccines during PHEs while maintaining the highest scientific and ethical standards. The Blueprint priority diseases, selected for their likelihood to cause PHEs and the lack of adequate medical countermeasures, were used to frame our methodological discussions. Here, we outline major vaccine study designs to be used in PHEs and summarize high-level recommendations for their use in this setting. We recognize that the epidemiology and transmission dynamics of the Blueprint priority diseases may be highly uncertain and that the unique characteristics of the vaccines and outbreak settings may affect our study design. To address these challenges, our group underscores the need for novel, flexible, and responsive trial designs. We conclude that assignment to study groups using randomization is a key principle underlying rigorous study design and should be utilized except in exceptional circumstances. Advance planning for vaccine trial designs is critical for rapid and effective response to a PHE and to advance knowledge to address and mitigate future PHEs.

**One Sentence Summary:** As part of the WHO research and development Blueprint for action to prevent epidemics, we describe key considerations for the design and analysis of trials and studies to evaluate experimental vaccines during public health emergencies.

## Introduction

The recent Ebola and Zika public health emergencies (PHEs) have demonstrated that the global community was not prepared to evaluate vaccines in affected countries, despite several decades of research into vaccine development on emerging pathogens *(1).* Epidemics of pathogens with no licensed vaccine will undoubtedly emerge in the future, and the public health community must be prepared to rapidly evaluate experimental vaccines in such circumstances. Our group was convened by WHO under the R&D Blueprint Plan of Action *(2)* with the mission to develop a consensus on study designs to rapidly evaluate vaccine candidates that address scientific, ethical and logistical issues arising in PHEs. We used the Blueprint priority diseases *(3)* to frame our discussions, to illustrate our rationale on key methodological considerations, and to anticipate future challenges.

The main goal of a vaccine efficacy trial is to obtain effectiveness data that can support broader use of a vaccine under a defined regulatory framework. In the context of an outbreak, vaccine evaluation also provides a way to give access, in the affected communities, to the most promising experimental vaccines and potentially to help control the current outbreak should the vaccine prove to be effective. In this process, we need to ensure that the experimental vaccine is demonstrated to be safe and effective and that it is used with an adequate community engagement and delivery strategy.

Conducting vaccine evaluation in PHEs is associated with methodological and operational challenges *(4, 5).* The epidemiology of the infectious disease, technological aspects, sociocultural aspects, and outbreak circumstances affect the choices we make when designing a vaccine trial or study.

We generally have limited knowledge about the transmission dynamics and the natural history of the Blueprint priority diseases. These pathogens are prone to epidemics where the spatiotemporal incidence of the disease may be highly variable and unpredictable. Unlike endemic diseases, outbreaks end or are contained to a point such that only sporadic cases occur. Furthermore, outbreaks may typically last only a few weeks and it may take one to two weeks for an outbreak to be detected and confirmed. These epidemiological and operational aspects make it difficult for studies to identify, enroll, and vaccinate at-risk participants prior to exposure, as well as defining the appropriate endpoints to estimate vaccine efficacy and effectiveness.

Given the sense of urgency that may arise, very little may also be known about the vaccine candidate itself in terms of safety and immunogenicity in humans, but also in terms of thermostability and other properties. Importantly, vaccine evaluation may also take place in a setting with unvalidated and unstandardized diagnostics and/or serologic assays, which poses considerable challenges for case ascertainment and endpoint measurement.

Outbreak circumstances are complex, and each outbreak has different characteristics. Typically, a PHE may trigger the rapid development of a large number of vaccine candidates that could be tested in affected countries. As a result, trial sponsors may compete for study sites and populations. In addition, research in epidemic management is relatively new. The conduct of research needs to be fully integrated into the international effort to control the disease, and should not be performed at the expense of the broader response to a PHE. Finally, there may be fears and misconceptions among the affected communities. Involving communities in the study implementation and complying with good participatory practices for research *(6)* are essential to increase acceptability of the intervention and preserve the integrity of the trial.

Because of the epidemiological situation and outbreak-working environment, we may not be able to conduct the perfect study. To address those challenges our group underscores the need for innovative, responsive and flexible study designs. As part of the Blueprint working group on vaccine evaluation, we present a summary of major vaccine study designs and design elements to be considered during PHEs of emerging and re-emerging pathogens for which there is no licensed vaccine.

## Results

It is widely acknowledged that double-blind placebo-controlled, individually-randomized vaccine trials performed in a variety of sites and study populations provide robust evidence that may contribute to inform licensure and broader use of a vaccine. However, in special circumstances, in the context of PHEs, trialists may be compelled to consider alternative study designs. Here, we outline major study design elements and challenges that are specific to the Blueprint priority diseases and to the context of PHEs, and we illustrate some of the trade-offs and methodological options.

### 1. Study endpoints

#### The challenges

For a given pathogen, study endpoints should be selected to support the broader intended use of a vaccine, as described in the WHO vaccine target product profile (TPP) for a given pathogen (Table **1**), and that is representative of the public health burden caused by that particular pathogen. However, it may not be feasible to have sufficient vaccine trial sample sizes with endpoints that are representative of the public health burden. In addition, if there are poor or limited diagnostics or limited infrastructure, endpoints requiring laboratory confirmation may be hard to detect. A clinical disease endpoint without laboratory confirmation should only be considered for pathogens with a highly distinct clinical syndrome, and these studies should consider laboratory testing of a random sample of cases to internally estimate the sensitivity and specificity of the case definition *(7).*

For instance, although cases of microcephaly represent the major public health burden associated with Zika infection, the choice of more frequent clinical events as a primary outcome measure for vaccine efficacy trials is likely to be necessary for feasible sample sizes *(8).* The justification of a mild, more common, endpoint as the primary endpoint in vaccine trials would be predicated on the assumption that the benefit of the vaccine on the selected endpoint is reasonably likely to predict clinical benefit for cases of microcephaly or other severe complications. Selecting an endpoint related to Zika infection would rely on a robust laboratory capacity and active surveillance system. However, there currently are no licensed diagnostics for Zika infection and serologic assays are cross-reactive with other arboviruses.

#### The methodological options

Methodological options include clinical endpoints or immunological surrogates of protection. Clinical endpoints, whether related to infection, disease or other clinical manifestations, may be clinically or laboratory confirmed. For instance, virologically-confirmed Zika illness is a convenient and feasible primary endpoint for a Zika vaccine efficacy trial *(8).*

#### Take home message

The demonstration of the vaccine benefit based on a clinical endpoint is the optimal way to evaluate a vaccine, but other approaches may be necessary if this is not feasible. Study endpoints may differ from the vaccine TPP desired from a public health perspective, but the benefit must be validated in future studies.

**Table 1.** Example of preferred characteristics extracted from the WHO vaccine Target Product Profiles *(9)* for Ebola, Lassa, MERS-CoV, Nipah and Zika, some of the pathogens prioritized by WHO.

|  | Indication for Use | Preferred measure of efficacy | Preferred target population |
| --- | --- | --- | --- |
| Ebola | For immunization of at-risk persons residing in the area of an on-going outbreak to protect against Ebola virus disease caused by circulating species of filovirus; to be used in conjunction with other control measures to curtail or end an outbreak | Preventing disease in healthy adults, adolescents and children | All age-groups and populations at high present risk of Ebola virus disease caused by circulating species of filovirus |
| Lassa | For active immunization of persons considered potentially at-risk, based on specific risk factors, to protect against Lassa disease. | Preventing infection or disease | All age groups.<br>Suitable for administration to pregnant women. |
| MERS-CoV | For active immunization of persons considered at-risk based on specific risk factors to protect against MERS-CoV. Risk groups will include health care workers, frontline workers and those working with potentially infected animals. | Preventing Middle East Respiratory Syndrome caused by MERS-CoV in healthy adults. | Health care workers, frontline workers and others with occupational risk.<br>Suitable for administration to pregnant women. |
| Nipah | For active immunization of at-risk persons in the area of an on-going outbreak for the prevention of Nipah disease; to be used in conjunction with other control measures to curtail or end an outbreak | Preventing Nipah infection or disease in healthy adults | All age groups and populations at high risk of Nipah disease |
| Zika | For the prevention of Zika virus-associated clinical illness of any severity in subjects 9 years of age or older | Prevention of virologically-confirmed Zika illness | Women of reproductive age (including adolescent and pre-adolescent girls 9 years of age or older), and boys/men of the same ages. |

### 2. Target population

#### Target population The challenges

Trial target population should also be representative of the target population defined in the vaccine TPPs (Table **1**). Likewise, it may not be feasible to have a sufficient sample size for a study population that is representative of the public health burden, for example, prevention of virologically-confirmed Zika illness in women of reproductive age *(8).* Typically, trials are implemented in sites with established high rates of disease transmission and draw participants from a general population representative of those who would ultimately receive the vaccine. However, because the incidence of new cases is extremely variable in PHEs, it may be challenging to identify a population in a given area that is at-risk and fully susceptible to disease transmission.

#### The methodological options

Study populations can be drawn from the general population or contact-based, or based on geographical or behavioral criteria. Studies may narrow the target population to those with other risk factors that make them at highest risk of infection, such as occupation. A targeted approach may require a smaller overall sample size if the incidence is truly higher in these individuals, though it may be harder to identify and to enroll these participants than a general population.

To take into account the high variability in disease transmission during an epidemic, we define a responsive target population as a study population that is triggered by the occurrence of a new case. In this regard, a responsive target population is designed to track the epidemic and focuses the intervention where the risk goes. For instance, the study population enrolled in the Ebola ring vaccination trial in Guinea *(10)* was a responsive contact-based study population where identification and enrollment of study participants was triggered by a confirmed case. This approach relies on a sensitive and rapidly responding surveillance system to inform the study in real-time as well as on a mobile and flexible vaccine delivery and cold chain. Such a design works best for single-dose vaccines that evoke a rapid immune response and for infectious diseases that spread relatively slowly through predictable contact networks.

**The take home message** – Responsive trials are appropriate in the event of an epidemic where the transmission dynamics are extremely variable in space and time, because they focus the intervention where the transmission and risk exposure are occurring, thereby increasing statistical power and decreasing the needed sample size.

### 3. Randomization

#### The challenges

Randomization provides assurance that the groups being compared are similar except for the vaccine being studied. The use of randomization was strongly debated in the context of the West-African Ebola outbreak *(11–13)* because the use of randomization may deny persons an opportunity to have access to a potentially effective vaccine in a situation with high mortality and lack of adequate medical countermeasures. Groups of experts argued that randomized trials are the most reliable and rapid way to identify the relative benefits and risks of investigational products and that every effort should be made to implement designs with random group assignment during outbreaks and epidemics *(12, 14).* Our group concurs with the above statement. Randomized trials are the study design of choice in PHEs, and deviation from the use of randomized designs should occur only under very exceptional circumstances under a robust risk-benefit analysis. For instance, if there is sufficient evidence of the safety and effectiveness of an investigational vaccine and there is no satisfactory alternative, the use of randomization may raise ethical concerns and acceptability among the affected populations.

#### The methodological options

A schematic of the different forms of appropriate randomized vaccine trials is shown in **Figure 1**. The unit of randomization can be at the individual or cluster level with various levels of stratification as pertinent.

**Figure 1:**
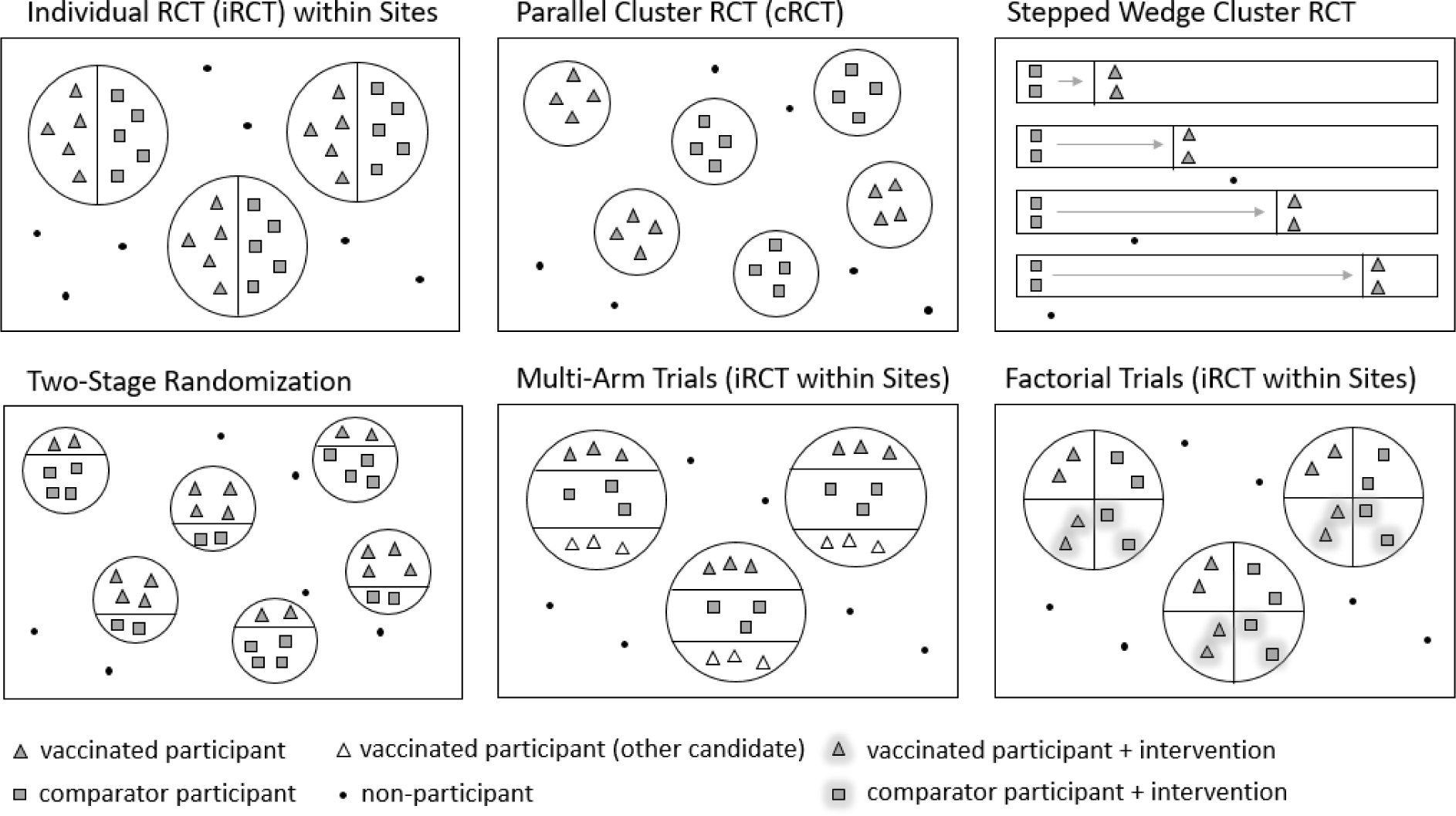
Schematic of randomized trial designs

#### Randomization at the individual level

Individually randomized controlled trials (iRCTs) can be customized for a PHE setting through creative definition of the study sites. Sites could be natural groupings of people at high risk of infection, or sites could be defined responsively following local disease spread. The primary analysis of an iRCT estimates the individual-level reduction in susceptibility to disease or infection (“direct vaccine effect” or sometimes “vaccine efficacy”). Population-level effects of vaccination, including indirect (spillover) protection, are typically not estimable *(15).* In some settings, high levels of indirect vaccine protection could dramatically reduce or shut off transmission in the comparator arm within small units *(16).* Though advantageous from a public health perspective, it would not be possible to evaluate vaccine efficacy (VE) in this setting. Finally, from an acceptability standpoint, participants may not consent to vaccinate some but not others within a small unit such as a household *(17).*

Multi-arm trials, where more than on vaccine is tested against a single placebo or comparator vaccine, are expected to require fewer resources than multiple, independent two-arm trials *(18).* Multi-arm trials allow direct comparison between the candidates because VE is measured concurrently in the same population using the same endpoints and methodology. This approach works best when the vaccines have complementary mechanisms of action and similar target populations. However, early evidence of efficacy in one candidate may prohibit further evaluation of other candidate(s) *(4).* From a public health perspective, this approach is attractive because it provides a method to evaluate concurrently a possibly large number of vaccine candidates evoked by the appearance of a PHE, has the potential to diversify the number and supply of vaccines available, and helps avoid monopoly situations in the long-run. This approach has been determined to be optimal for Zika vaccine trials where future transmission will probably occur in different geographic clusters in pockets of still susceptible populations.

In the absence of effect modification, factorial trials (iRCTs within sites factoring in nonvaccine interventions) have the same power as two independent trials, and conserve resources by utilizing the same population and trial infrastructure *(19).* This approach may be considered when the vaccine and other experimental interventions *(e.g.*, vector control methods) have overlapping eligibility criteria, complementary mechanisms of action, and similar toxicity profiles. In the case of vector-borne diseases, this approach is attractive because it provides a way to reconcile innovative vector control evaluation and vaccine evaluation against the same disease in a given area.

#### Randomization at the cluster level

Parallel cluster randomized controlled trials (parallel cRCTs) are well-suited for infectious diseases that exhibit transmission in clustered populations, such as the Blueprint priority diseases, because clusters can capture transmission networks *(17, 20).* Clusters should not overlap to reduce contamination *(17).* For PHEs, clusters provide programmatic advantages and can be defined to capture high-risk individuals or responsively follow local transmission. However, parallel cRCTs may be difficult to blind and are thereby subject to a number of biases that can reduce interpretability of the results *(21).* The primary analysis estimates total vaccine effectiveness, which combines direct and indirect vaccine effects *(15)*, and, if data collection were expanded to include non-participants, the trial could produce estimates of indirect and overall effects. A form of this design was used for the successful Ebola ring vaccination trial in Guinea *(22).*

In the context of PHEs, stepped wedge cRCTs have important disadvantages, primarily because they are complex to plan, implement, and analyze *(17).* Stepped wedge cRCTs are inflexible, as all participants and facilities must be enrolled before the first dose of vaccine can be administered, even though vaccine is delivered in a time-staggered fashion (23). Stepped wedge cRCTs probably result in the slowest trials and are not well-suited for endpoints with spatiotemporally variable incidence *(23–25).*

Two-stage randomized designs are one of the only designs to support relatively unbiased estimation of both direct and indirect vaccine effects *(26).* An important disadvantage of the design is its complexity, and there is no precedent for such design in vaccine trials.

**The take home message** – Despite the exceptional circumstances of a PHE, randomization, whether at the individual or cluster level, remains a key principle in vaccine evaluation. Deviation from the use of randomized designs should occur only under very exceptional circumstances. For PHEs, we recommend randomized trial designs that are compatible with the enrollment of a responsive target population. cRCTs can provide both direct measures of vaccine efficacy, as well as measures of indirect vaccine effectiveness, while iRCTs only provide measures of direct effectiveness *(15).*

### 4. Comparator

#### The challenges

A common model for evaluating and deploying a new vaccine, against a disease for which there is no existing vaccine, is that it is first tested in a trial controlled with a placebo or with an unrelated vaccine. The use of blinding (or masking), as is possible with the use of a control, reduces the potential for biases, such as selection, detection, and performance bias *(27).* Like with randomization, the use of a placebo has been strongly debated in the context of the West-African Ebola outbreak *(28)* and will likely be debated in future PHEs. In contrast, a delayed vaccination approach used as a comparator arm was implemented in the Ebola ring vaccination trial in Guinea *(22, 29)*, and the Ebola iRCT trial in Sierra Leone *(30).*

#### The methodological options

Researchers should consider whether the risks associated with use of the placebo – that is the risks of the placebo intervention itself and those of withholding or delaying a vaccine with demonstrated efficacy and effectiveness – are minimal, preventable or reversible. Risks greater than this may constrain the use of placebos.

In PHEs, a delayed vaccination comparator may be adopted in which individuals/clusters are allocated to either immediate or delayed vaccination. Motivations for the use of a delayed comparator include improving acceptability, providing vaccine to individuals in greatest need, and averting more cases and promoting epidemic control if the vaccine is efficacious. However, if the vaccine is ineffective or dangerous, more people are exposed to the vaccine than would be in a trial with placebo or unrelated vaccine control. This approach is expected to have lower power and the outcome estimates may be biased *(31).* To reduce bias, the length of the delay should be relatively long compared to the disease incubation period and the time required for the immune response to develop among vaccinated people.

In settings where an existing vaccine has already been established to provide clinically meaningful benefit, an experimental vaccine may have potential advantages other than efficacy, such as having a more favorable tolerability or safety profile, being more convenient to store, transport, or administer, or less costly. It might be sufficient for the experimental vaccine to have similar rather than superior efficacy relative to the existing vaccine, which can be evaluated in a non-inferiority trial *(32).* Depending on the size of the non-inferiority margin (minimum threshold for an unacceptable loss of efficacy), non-inferiority trials may require large sample sizes that make them challenging in the PHE setting.

#### The take home message

Although the use of placebo provides a robust methodological standard, the use of delayed vaccination can be explored in certain circumstances.

### 5. Primary analysis

The primary analysis can be conducted per protocol, intention-to-treat (ITT), or modified ITT. The per protocol analysis restricts the population to eligible, fully compliant participants receiving all doses as allocated per protocol. The analysis includes a delay, usually starting after the final dose of the vaccine plus the maximum incubation period, to allow the immune response to develop and to account for the time between infection and symptom onset. The goal of the per protocol analysis is to estimate the intrinsic efficacy of the vaccine to support licensure decisions and planning, but it is subject to post-randomization biases such as differential loss to follow-up. Alternatively, an ITT analysis includes all cases occurring after randomization or all cases occurring after the first dose of vaccine/placebo. The ITT analysis yields a practical, though more context-specific, estimate of vaccine effectiveness because it includes cases who may have been infected before the vaccine induced an immune response, as well as individuals who fail to comply with the protocol, potentially for reasons relating to the vaccine itself. As a result, the ITT estimate of VE tends to be attenuated compared to the per protocol estimate, and the difference between the ITT and per protocol estimates of VE may be especially large if many infections occur during the per protocol delay *(31).* In the modified ITT approach, a sensitive test is used to retrospectively exclude individuals infected at baseline *(33)*, though this requires the availability of both baseline samples and a reliable test. Although ITT is the preferred approach in clinical trials, VE trials typically conduct a per protocol primary analysis *(34).* Though only a single primary analysis may be selected, both ITT and per protocol estimates of VE should be reported.

## Discussion

In this document, we have outlined major study designs and design elements to be considered in PHEs. A key principle is that randomized designs should be utilized whenever possible. Observational studies *(e.g.*, cohort studies, case-control and test-negative designs *(15, 35, 36))* are non-randomized and therefore should only be considered in certain limited settings because the quality of inference will always be viewed as inferior relative to a randomized design. A setting where observational studies may be useful is when the product of interest has received conditional licensure but needs to be further evaluated. Like any observational study, collection of and adjustment for potential confounders is critical. Results of observational studies are easiest to interpret when the effect of the intervention is large enough so as to overshadow random error and bias, especially due to confounding *(37).* Furthermore, human challenge studies, where participants are intentionally exposed to the pathogen, can use classical experimental designs and relatively small sample sizes to directly assess efficacy, safety, and immunogenicity of an experimental vaccine. In rare settings, human challenge studies may be used to support regulatory decisions, provided that that the human challenge model system is adequately predictive of efficacy in the field *(38).*

For randomized trials, independent Data and Safety Monitoring Committees should be in place to safeguard the interests of study participants and to enhance the integrity and credibility of the vaccine trial *(39).* The design of the vaccine trial should include specification of data monitoring boundaries allowing for early termination of the trial for benefit or for futility while controlling the trial’s type 1 error rate and preserving power. Group sequential guidelines, such as an O’Brien-Fleming or Pocock boundary, provide a widely implemented approach *(40, 41).* The number and timing of interim analyses can be flexibly defined through an alpha spending method *(42).* The flexibility of interim analyses is important because it may allow a trial to be stopped at the earliest opportunity in order to discard a futile or unsafe vaccine or to further endorse an experimental vaccine to influence the course of the current outbreak. In this regard, the ability of a trial design to be transformed into an intervention that can influence the course of the outbreak is also essential. Following the promising results of the rVSV-ZEBOV vaccine against Ebola virus disease *(29)*, the targeted and flexible nature of the ring vaccination approach used in Guinea allowed for the rapid transformation of rings into an intervention in response to a flare-up of Ebola transmission, several months after West Africa was declared Ebola-free, and where the vaccine was deployed under compassionate use criteria *(43).* If a trial is terminated early, there should also be a plan in the protocol for next steps, which may include vaccinating all eligible, consenting, unvaccinated participants. These participants should then be followed for safety outcomes since the product would be unlicensed at that time.

Adequate guidance does not exist on how to conduct data monitoring in the presence of changing epidemiologic conditions. For example, transmission among humans may decline to extremely low levels or stop entirely, precluding accrual of further evidence to directly evaluate vaccine efficacy. The protocol should clarify how study data would be analyzed in this scenario, including alpha spending, as the fully needed sample size would not have been reached. A waning epidemic could trigger study closure with a final analysis, study pause until the next outbreak occurs in that area, or study continuation to collect additional safety and immunogenicity data. Keeping the study open would also be desirable in case there is an unexpected surge in transmission. This decision could be guided by an evaluation including transmission modelling to assess the probability of future cases in the current outbreak or future outbreaks in the study area. This type of modelling has been used to inform likely case accrual for Ebola vaccine trials *(44).* It may be difficult to accumulate enough evidence to reliably ascertain the efficacy of an intervention from a single outbreak. As a result, it may be necessary to proactively plan to combine information across multiple outbreaks (or trials within the same outbreak) to evaluate the efficacy of an intervention. The methods for combining evidence from randomized trials across outbreaks are covered in reference *(45).*

Given the circumstances of PHEs and the epidemiological situation, we have underscored the need for responsive and flexible study designs while maintaining the highest scientific and ethical standards possible. An interactive, web-based decision support tool has been developed to navigate through the various study design elements and options outlined here and to promote scientific discussion among methodologists *(46).* Our group also recognizes the role of mathematical models of infectious disease to explore different assumptions and analyze their impact on the statistical power of a given vaccine trial design in a given epidemic scenario *(47, 48).*

Many of the principles described here for vaccine studies can be expanded to therapeutic and prophylactic antimicrobial agents. By expanding these study designs and plans for all potential emerging infectious disease threats on the Blueprint priority disease list, we will be able to rigorously evaluate vaccine and antimicrobial efficacy and effectiveness at the earliest opportunity when an outbreak occurs, to mitigate current and future outbreaks.

## Acknowledgments

The authors are members of the WHO R&D Blueprint workplan for designing clinical trials in Public Health Emergencies, and have participated in a series of consultations. The authors thank WHO and the other members of the Blueprint workplan,including Thomas Fleming, Susan Ellenberg, Philip Krause, Jonathan Sterne and Yazdan Yazdanpanah for their thoughtful feedback. Funding: NIH R37-AI032042 (NED, IML, MEH), NIH U54-GM111274 (NED, IML, MEH): NED, IML and PSG drafted the manuscript. All authors reviewed and edited the manuscript. Competing interests: The authors declare no conflict of interest.

